# Engineered phenotype patterns in microbial populations

**DOI:** 10.1101/575068

**Authors:** Philip Bittihn, Andriy Didovyk, Lev S. Tsimring, Jeff Hasty

## Abstract

Rapid advances in cellular engineering^1,2^ have positioned synthetic biology to address therapeutic^3,4^ and industrial^5^ problems, but a significant obstacle is the myriad of unanticipated cellular responses in heterogeneous environments such as the gut^6,7^, solid tumors^8,9^, bioreactors^10^ or soil^11^. Complex interactions between the environment and cells often arise through non-uniform nutrient availability, which can generate *bidirectional* coupling as cells both adjust to and modify their local environment through different growth phenotypes across a colony.^12,13^ While spatial sensing^14^ and gene expression patterns^15–17^ have been explored under homogeneous conditions, the mutual interaction between gene circuits, growth phenotype, and the environment remains a challenge for synthetic biology. Here, we design gene circuits which sense and control spatiotemporal phenotype patterns in a model system of heterogeneous microcolonies containing both growing and dormant bacteria. We implement pattern control by coupling different downstream modules to a tunable sensor module that leverages *E. coli⁉s* stress response and is activated upon growth arrest. One is an actuator module that slows growth and thereby creates an environmental negative feedback via nutrient diffusion. We build a computational model of this system to understand the interplay between gene regulation, population dynamics, and chemical transport, which predicts oscillations in both growth and gene expression. Experimentally, this circuit indeed generates robust cycling between growth and dormancy in the interior of the colony. We also use the stress sensor to drive an inducible gating module that enables selective gene expression in non-dividing cells. The ‘stress-gated lysis circuit’ derived from this module radically alters the growth pattern through elimination of the dormant phenotype upon a chemical cue. Our results establish a strategy to leverage and control the presence of distinct microbial growth phenotypes for synthetic biology applications in complex environments.

---

To investigate bacterial populations under spatially-inhomogeneous conditions, we built a microfluidic device based on our previous design^18^, where deep side-traps protrude from nutrient supply channels that continuously deliver fresh media to one side of the growing colony while washing away spillover cells (Fig. 1a; Supp. Figs. 1 and 2). When loaded into the device, E. *coli* colonies grew to fill the traps. Cells deprived of nutrients at the rear of the traps switched to a nondividing phenotype resembling the small, round morphology in stationary-phase batch cultures^19^ (Fig. 1b; Supp. Figs. 3, 4; Supplementary Movie 1), indicating that nutrient supply is inherently diffusion-limited^20^. In the rich medium used here (see Methods), the steady-state growth pattern featured a sharp boundary (at ~2 hours) between rod-shaped, dividing and small non-dividing cells (Fig. 1c).

**Figure 1.**
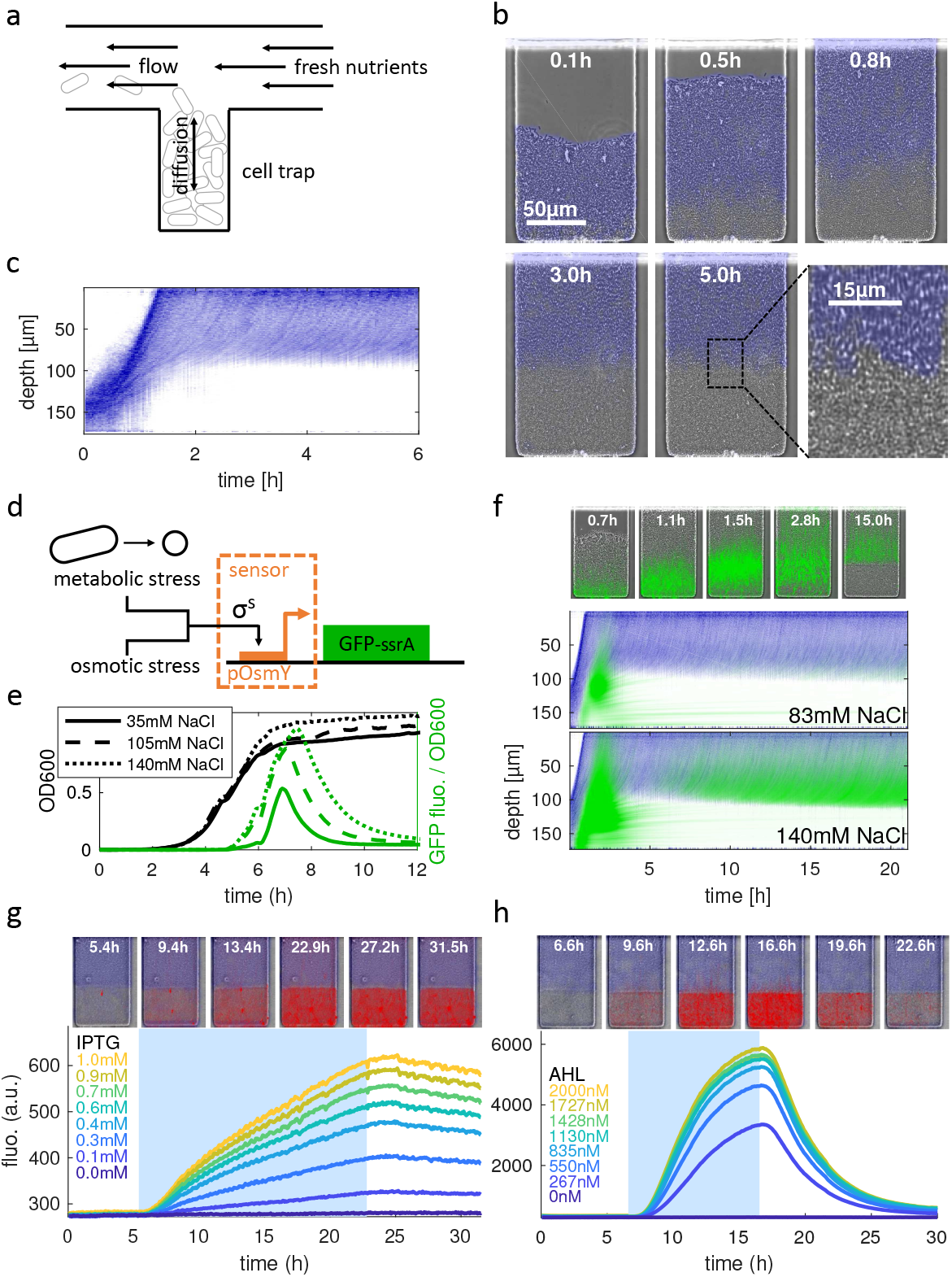
Microfluidic platform for observing *E. coli* growth patterns in spatially heterogeneous microcolonies. **a**, Schematic illustrating a trap in our microfluidic device. The flow rate in the nutrient supply channel is ≈100*μ*m/s. **b**, Time-lapse microscopic images of a single microfluidic trap showing development of phenotypic heterogeneity shortly after cell loading (see Supplementary Movie 1). Blue color indicates regions of actively growing *E. coli* (see Methods). **c**, Kymograph of image differences (blue) averaged along the horizontal dimension of the trap. **d**, The stress sensor consists of the σ^S^-responsive pOsmY promoter driving ssrA-tagged GFP to visualize sensor activation (panels e and f). **e**, Time courses of OD and fluorescence in plate reader experiments display transient sensor activation that depends on the level of basal osmotic stress (NaCl). **f**, Time-lapse images (for 140 mM NaCl; see Supplementary Movie 2) and kymographs of microfluidic experiments showing NaCl-dependent transient stress sensor activation (green) upon initial growth arrest and later persistent activation in dividing cells close to the no-growth boundary. **g**, Protein expression in growth-arrested *E. coli* measured by induction of IPTG-inducible pLlacO-1 driving RFP In the time-lapse images, red indicates RFP fluorescence, blue marks regions of growth. Plotted time traces represent average RFP fluorescence in the growth-arrested region of the trap for different IPTG concentrations present during the induction window (blue shaded area). Expression in growth-arrested cells was also observed with pLuxI and AHL (Supp. Fig. 7). **h**, The same for pLuxl-driven RFP which is ssrA tagged and targeted for enzymatic degradation by ClpXP.

**Figure 2.**
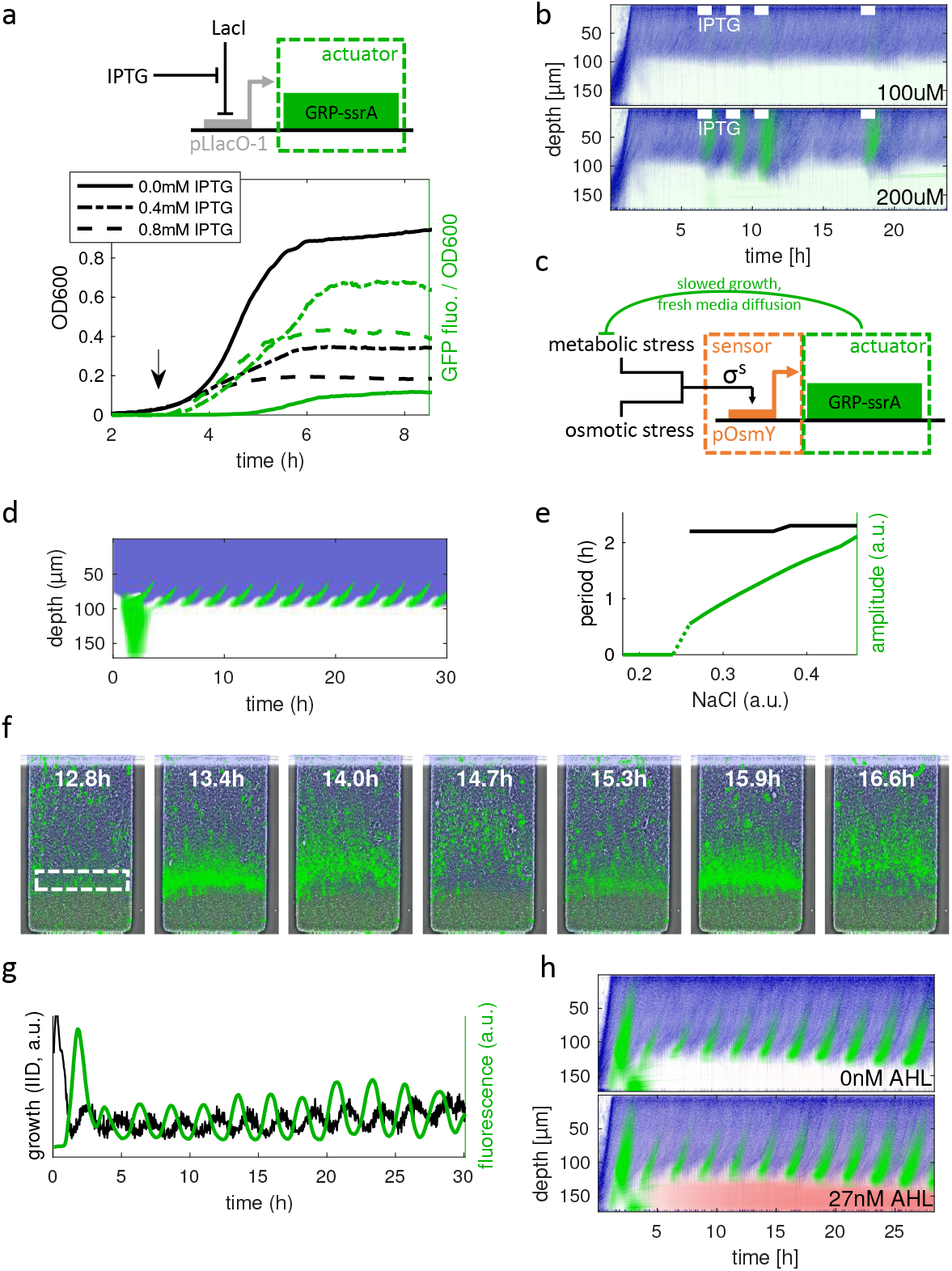
Actuator growth modulation and spatiotemporal feedback. **a**, Schematic of the actuator circuit expressing GRP-ssrA from the pLlacO-1 promoter. The circuit was induced with IPTG at OD 0.03 (black arrow, see Supp. Fig. 8a for induction at OD 0.13). **b** Kymographs showing growth (blue) and GFP fluorescence (green) of *E. coli* cells with the actuator circuit in microfluidics. Pulses of IPTG shown by white marks. (See Supp. Fig. 8 for time-lapse microscopic images and behavior across all tested IPTG concentrations, and Supp. Fig. 9 for a continuous induction of the actuator circuit.) **c**, Sensor-actuator circuit. The pOsmY stress sensor drives the GRP-ssrA actuator protein, giving rise to spatiotemporal negative feedback mediated by growth suppression that increases available nutrient concentration and relieves metabolic stress. **d**, Kymograph from numerical simulations of the spatiotemporal model of microcolony dynamics with the sensor-actuator circuit. After the sensor and actuator components were separately tuned to match the experimental dynamics observed in batch and microfluidic experiments with isolated circuits (Figs. 1e, 2a, b), periodic oscillations of actuator protein concentration (green) and growth (blue) were observed in simulations of the full model near the no-growth boundary (see Supp. Fig. 10 and Supplementary Information 3). **e**, Period and amplitude of actuator protein concentration oscillations in the spatiotemporal simulations. **f**, Time-lapse images of cells carrying the sensor-actuator circuit showing oscillations of GRP-ssrA fluorescence (green) at the growth boundary in media supplemented with 71mM NaCl needed for feedback activation (cf. Supp. Fig. 11; see Supplementary Movie 4). **g**, Local oscillations in GRP-ssrA fluorescence and growth (calculated via Inter-image differences, see Methods) averaged over the area marked in panel f. **h**, Kymographs showing periodic oscillations of GRP-ssrA fluorescence (gree) and growth (blue) in a microfluidic colony of cells with the sensor-actuator circuit and expressing untagged RFP (red) from an AHL-inducible pLuxI promoter.

As the first step toward engineering growth-state coupled regulatory circuits, we constructed a ‘stress sensor module’ which is activated upon transition into stationary phase, when the alternative/stationary phase sigma factor (σ^S^) becomes active. Our sensor is based on the σ^S^-dependent promoter pOsmY (see Methods) which, in addition to responding to metabolic stress (nutrient deprivation), is tunable by NaCl concentration^21^. When it was used to drive GFP expression (Fig. 1d), we observed a sharp, osmolarity-dependent increase in fluorescence upon entry into stationary phase in batch culture (Fig. 1e). Sensor activation in microfluidics occurred upon initial microcolony growth arrest and in dividing cells close to the no-growth boundary when exposed to higher osmolarity, consistent with batch culture results (Fig. 1f; Supp. Fig. 6; Supplementary Movie 2).

To monitor protein expression we transformed the cells with a construct expressing RFP from a LacI-repressed pLlacO-1 promoter^22^. IPTG-dependent RFP expression was observed for more than 15 hours in dormant cells (Fig. 1g). Similar results were obtained for the pLuxI promoter upon induction with N-(3-oxohexanoyl)-homoserine lactone (AHL; Supp. Fig. 7). consistent with long-term gene expression previously seen in stationary cells^23^. For both promoters’ the decline of fluorescence upon cessation of induction was minimal due to the absence of dilution by cell growth. We tested whether enzymatic degradation by the unfoldase-protease ClpXP^24^ was also long-lived in dormant cells. Targeting RFP for degradation by adding an ssrA degradation tag led to a rapid decline of fluorescence upon removal of the inducer (Fig. 1h), indicating that non-dividing cells support both long-term expression and degradation *in situ*.

Next we looked for a way to actively modulate the phenotype pattern within the microcolonies. We wondered whether non-dividing cells contributed to the chemical conditions that caused growth arrest and affected the position of the no-growth boundary. Comparing traps with depths of 170*μ*m and 140*μ*m—the latter containing about half as many non-dividing cells as the former—we observed very similar growth patterns relative to the mouth of the trap (Supp. Fig. 5a, b), indicating that non-dividing cells do not actively participate in growth pattern formation. Therefore we focused our attention on modulating cell growth of the dividing cells.

As an actuator module, we constructed a fluorescent growth repression protein (GRP) by fusing a portion of the OsmY protein with green fluorescent protein (see Methods) and ssrA-tagged it to eliminate toxicity at basal expression levels. When expressed from an IPTG-inducible promoter GRP-ssrA substantially reduced and ultimately stopped cell growth (Fig. 2a). In the microcolonies growing within microfluidic traps, growth rate reduction of dividing cells allowed nutrients to diffuse deeper into the traps. Pulsed expression of GRP-ssrA reversibly modulated the phenotype pattern by slowing growth in the front of the trap and causing previously dormant cells in the back to temporarily resume growth (Fig. 2b; Supplementary Movie 3).

We combined the actuator module with our tunable stress sensor module (Fig. 1d) to reduce growth in the cells experiencing metabolic stress, thereby facilitating nutrient diffusion into the trap (Fig. 2c). This circuit resembles a negative feedback loop, where a gene product suppresses its own production, promoting oscillatory behavior with the distinction that the feedback loop here is mediated by environmental diffusion. A computational reaction-diffusion-advection model based on mass-action kinetic equations for our sensor and actuator modules (Supp. Fig. 10) coupled with the equation for the nutrients (see Supplementary Information 3) predicted that oscillations should be observed when the stress sensor module is tuned to sufficiently high expression levels (Figs. 2d and e, Supp. Fig. 10d). Experimentally’ these oscillations were observed in a layer of cells close to the no-growth boundary as regular repetitive modulations of both GRP fluorescence and growth (Supplementary Movie 4, Figs. 2f and g) for sufficiently high NaCl concentrations (≳ 40mM). The dormant layer of cells behind the oscillatory region remained undisturbed and supported RFP expression (Fig. 2h). Consistent with the theoretical prediction (Fig. 2e; Supp. Fig. 10d), no oscillations were observed in the absence of NaCl (Supp. Fig. 11a), but, once induced, the period was insensitive to the sensor induction level determined by NaCl (Supp. Fig. 11b).

While our actuator module has the ability to increase the diffusive flow of nutrients to the back of the trap and thus reactivate some dormant cells (Fig. 2b), growth only recovers up to a certain depth beyond which cells remain in stationary phase. This phenotypic heterogeneity could be undesirable, as altered physiology of the bacterial host could compromise gene circuit function, and the absence of growth-mediated dilution could distort expression levels such as readouts from biosensors. Therefore, we devised an inducible gene circuit that would eliminate stationary phase cells and thus restore population homogeneity. We used phage ϕX174 lysis protein E, which had been used previously for population control^8, 25, 26^. However, expressing the lysis protein from the standard pLuxI promoter was not able to selectively eliminate dormant cells without severely affecting dividing cells (see Supplementary Information 2, Supp. Fig. 12 and Supplementary Movie 5).

To counter the problem of unintended lysis protein synthesis in dividing cells, we constructed a “gating module” designed to limit expression of a target gene specifically to growth-arrested cells. We removed the LuxR transcription factor, which activates the pLuxI promoter in the presence of AHL, from the standard pLuxI cassete and instead expressed it from the pOsmY stress sensor promoter (Fig. 3a), priming cells for pLuxI induction by AHL only upon growth arrest. In the presence of the gating module, LuxR and target gene levels should be reduced in dividing cells and increased in growth-arrested cells compared with constitutive LuxR expression in the standard ungated pLuxI promoter cassette (Fig. 3b).

**Figure 3.**
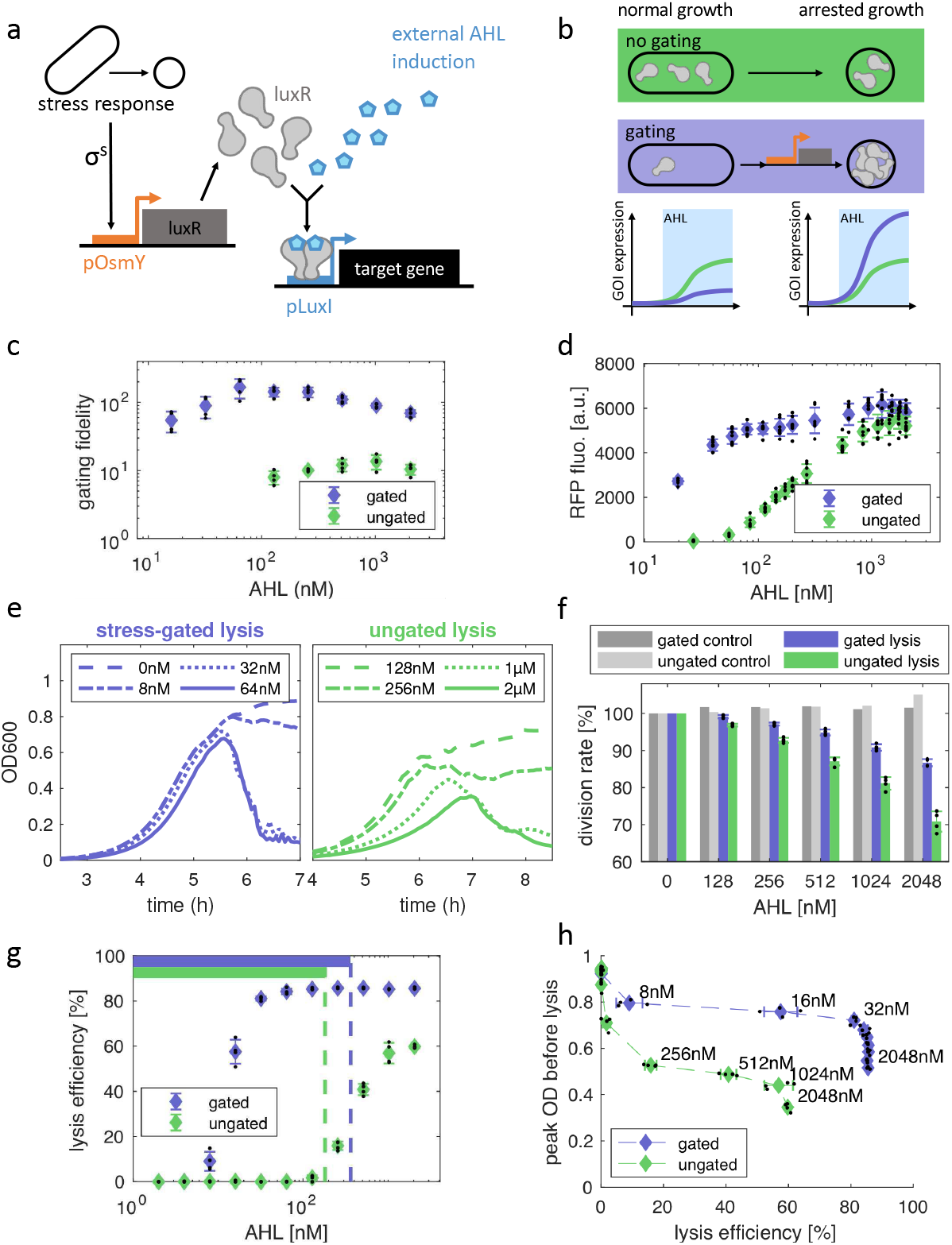
Synthetic stress gate primes cells for phenotype-specific expression. **a**, Stress-gated expression of a target gene, the pOsmY stress sensor drives expression of the regulator protein LuxR required to activate the pLuxI promoter upon binding of external AHL. **b**, Constitutive LuxR expression (green) from the native pLuxR promoter in the standard pLuxI induction cassette (cf. Supp. Fig. 17) leads to sensitivity of both growing and growth-arrested phenotypes to pLuxI induction by external AHL. The gating module (purple) reduces LuxR levels during normal growth and primes cells with LuxR upon transition into the non-growing phenotype, increasing pLuxI promoter sensitivity and thus target gene expression upon induction with external AHL in a phenotype-specific manner. **c**, Stress-gated RFP circuit (target gene = RFP-ssrA-AAV) with constant AHL induction in the plate reader compared to an ungated circuit. Gating fidelity calculated by dividing peak RFP fluorescence (cf. Supp. Fig. 13b) by RFP fluorescence at OD 0.7. Data shown is for peak fluorescence values above 5% of peak fluorescence at maximum induction. Error bars indicate standard deviation (SD) across 4 technical replicates. **d**, The same circuits cultured in microfluidics and induced for 10 h at different AHL levels. Average RFP fluorescence in the growth-arrested region was determined as in Fig. 1h. Error bars indicate SD across 7 microfluidic traps. **e**, Batch growth of cells with the stress-gated lysis circuit (panel a with target gene = E-ssrA-AAV) and the ungated circuit at various inducer concentrations. **f**, Growth rates for the stress-gated and ungated lysis circuit, and for corresponding controls expressing RFP-ssrA-AAV instead of E-ssrA-AAV. Error bars indicate SD across 4 technical replicates. **g**, Lysis efficiency (percentage OD reduction within 1h after reaching peak OD) for the two lysis circuits. Horizontal bars at the top of the plot indicate the inducer range where the growth rate reduction in exponential phase is less than 5%. Error bars indicate SD across 4 technical replicates. **h**, Peak OD reached before lysis vs. lysis efficiency for the gated and the ungated lysis circuit. Horizontal error bars indicate SD across 4 technical replicates. SDs of peak ODs are smaller than plot symbol sizes.

To test this design, we placed RFP as the target gene under the control of the gating module and tagged it with a weak ssrA degradation tag (“ssrA-AAV”; see Methods) to prevent accumulation from leaky expression in dormant cells. We defined “gating fidelity” as the ratio of the maximum fluorescence after the onset of stationary phase to the fluorescence at a specific low OD in batch culture (see Methods and Supp. Fig. 13a). We observed a greater than ten-fold increase in gating fidelity (Fig. 3c) compared to non-gated expression, together with a stronger response at much lower AHL levels both in batch culture (Supp. Fig. 13b) and in microfluidics (Fig. 3d). Thus, the gating module was effective in increasing expression in growth-arrested cells relative to dividing cells. This growth-state specificity was maintained across different growth media (Supp. Figs. 13d, e). By reducing LuxR expression during exponential growth, the gating module was also able to improve exponential-phase growth rate compared to constitutive LuxR expression (Supp. Fig. 14). In addition, induction time—the time to reach 20% of the final fluorescence— in non-dividing cells was substantially reduced compared to the standard (non-gated) divergent-expression pLuxI cassette. The induction time was constant across all AHL concentrations (Supp. Fig. 13c), in contrast to the strong dependence on the AHL concentration in the standard pLuxI cassette, where it was possibly caused by the positive autoregulation of LuxR^27^.

Placing lysis gene *E* tagged with the same ssrA-AAV degradation tag as the target gene under the control of the gating module we obtained a “stress-gated lysis circuit” (SGLC). We assessed the circuit’s behavior in batch culture by analyzing growth at different induction levels of AHL and comparing it to ungated expression of the lysis gene from the standard pLuxI expression cassette. Lysis for the SGLC occurred at lower AHL concentrations than for the ungated circuit, consistent with the results of the gated RFP test circuit. At AHL concentrations that were effective, the SGLC caused rapid onset of lysis after normal exponential growth, while initial exponential growth was slowed by the ungated circuit (Fig. 3e). To quantify these differences, we determined exponential growth rate at low OD, which was significantly less diminished in the gated lysis circuit compared to the ungated circuit (Fig. 3f). At the same time, the SGLC achieved high lysis efficiency—defined as the reduction in OD one hour after the peak—at much lower AHL concentrations, creating a regime not seen in the ungated circuit: A wide range of AHL concentrations caused strong lysis upon entry into stationary phase, but no detectable growth rate reduction in exponential phase (Fig. 3g, cf. Fig. 3e). Similarly, in the non-gated circuit, effective stationary-phase lysis was only observed at the cost of substantial peak-OD reduction, whereas, for the SGLC, lysis efficiency was generally high and the change in peak OD was small over more than two orders of magnitude of AHL concentrations (Fig. 3h), indicating effective “gating”.

In our diffusion-limited microcolonies, the SGLC achieved specific, long-term elimination of growth-arrested cells. In contrast to the ungated circuit, lysis occurred only in the dormant cells: No lysis was observed in the dividing subpopulation (Fig. 4a; Supplementary Movie 6). Instead, the steady-state growth pattern that developed superficially resembled the original population structure, with a sharp boundary (Fig. 4b) and dormant cells replaced by debris from previous lysis events. During continued exposure to sufficiently high AHL levels (≳ 9nM), any cells transitioning to the non-dividing phenotype in the region with inadequate nutrient supply were rapidly eliminated (Fig. 4c). AHL levels mainly determined the time until sufficient lysis protein E had accumulated for the onset of lysis (Fig. 4d; Supp. Fig. 15b; Fig. 4e). At low AHL concentrations, this increased time to lysis led to oscillations, characterized by repeated trap fillings alternating with lysis events several hours after growth arrest (Fig. 4d; Supp. Fig. 16; Supplementary Movie 7).

**Figure 4.**
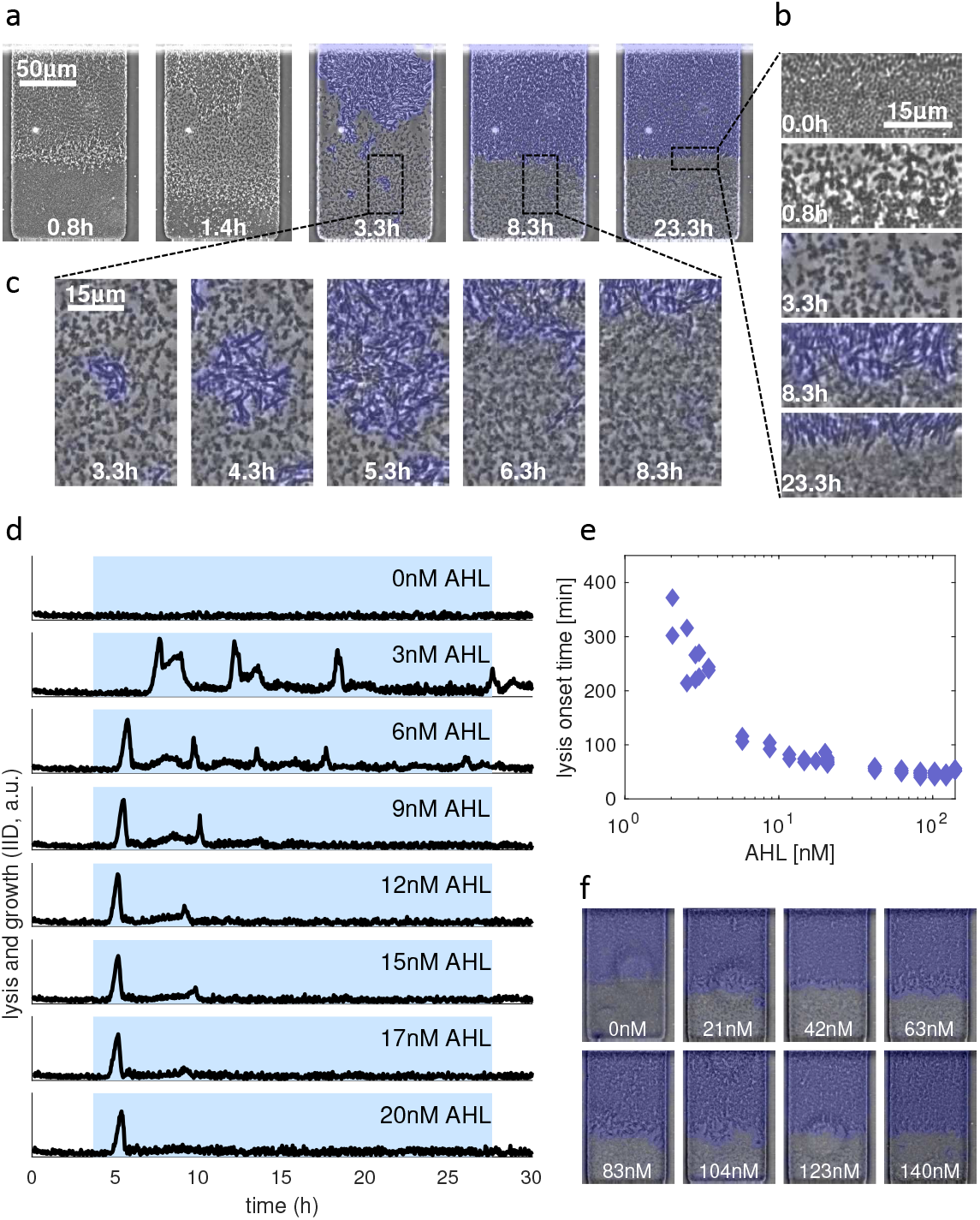
A colony-level phenotype filter. **a**, Time-lapse images of a single microfluidic trap with cells carrying the SGLC (see Supplementary Movie 6). Dormant cells are eliminated from the population while dividing cells are unaffected by the SGLC. Blue = regions of active growth after the first lysis event. Time stamps indicate time after beginning of 17nM AHL induction. **b**, Magnification of the interface region for the same time points as in panel a. **c**, Magnification of a region in the rear of the trap, showing a small population of cells growing after the first lysis event that eventually lyses when steady-state growth in the front part of the trap is established. **d**, Growth and lysis dynamics across AHL inducer levels for the SGLC. Curves were obtained by calculating inter-image differences (see Methods) in the region of the trap usually occupied by growth-arrested cells. Spikes correspond to sudden lysis events. Blue-shaded regions denote AHL induction windows. **e**, Time from induction to the onset of lysis was determined by detecting the first peak in the IID traces as in panel **d**. Experimental data was pooled from three different experiments for 0-3.5nM, 0-20nM and 0-140nM AHL. Each data point corresponds to a single trap on the microfluidic chip. **f**, Time-lapse images of the steady state taken after 18h of AHL induction with the indicated concentration.

Overall, very similar steady-state phenotype patterns, consisting only of dividing cells, developed over more than one order of magnitude of inducer levels (Fig. 4f; Supp. Fig. 15a), reflecting the performance of the gating circuit and the insensitivity of exponentially growing cells to the SGLC in batch over a wide range of AHL concentrations (cf. Fig. 3h).

The expanding synthetic biology toolkit permits the engineering of microbes with increasingly complex functionality^2, 28^. Our results represent an additional design layer, providing fine-grained control over internal population structure. Stress-gated lysis circuits could be used for targeted drug delivery^29^ triggered not by crossing a specific quorum-sensing threshold^8^, but by changes in cell phenotype. More generally, growth pattern engineering could provide methods to control the gut colonization behavior and microbiome interactions of synthetic probiotic bacteria^7^ and improve biocontainment strategies that prevent the undesired accumulation of dormant designer microbes in natural environments^30^. Furthermore, exploiting the coexistence of different phenotypes in conjunction with phenotype-gated circuits might also allow for engineering new types of division of labor among isogenic cells.

## Acknowledgements

The microfluidic mixer is derived from a design developed by Megan Dueck. We also thank Garrett Graham for stimulating discussions in the early phase of the project and Quoc Tran for technical help. This material is based upon work supported by the NIH/NIGMS grant RO1-GM069811 and NSF grant No. DMS-1463657. P.B. was supported by the Human Frontiers Science Program fellowship LT000840/2014-C. Any opinions, findings, and conclusions or recommendations expressed in this material are those of the authors(s) and do not necessarily reflect the views of the funding agencies.

## Author Information

The authors declare no competing financial interests. All authors (P.B., A.D., L.S.T., J.H.) contributed extensively to the work presented in this paper. Correspondence and requests for materials should be addressed to L.S.T. or J.H.

## Methods

### Plasmids and Strains

All DNA constructs were created using EMILI^31^ (for deletions and short modifications) or Gibson Assembly^32^ (for assembly of multiple parts). Templates for PCR were either previous constructs from our group^8,18,33^ (inducible promoters and fluorescent proteins) or the *E. coli* MG1655Z1 genome (stress promoters). To target proteins for degradation by the protease ClpXP, strong and weak ssrA degradation tags^34^ with amino acid sequences “AANDENYALAA” and “AANDENYAAAV”, respectively, were fused to the C-terminus of the corresponding protein, encoded immediately before the stop codon. All constructs were verified by Sanger sequencing. Addgene catalog numbers for plasmids will be provided upon publication.

Our initial studies of σ^S^-dependent promoters included those for genes *poxB, yciG* and *osmY* based on their previously shown strong response to multiple stresses^21^. We eventually settled on *osmY*, as it showed a particularly peaked response upon entry into stationary phase and tunability by NaCl (Fig. 1e). The essential part of our standard pOsmY promoter was PCRed from the genome as nucleotides −235 to 427 relative to the transcriptional start site of *osmY*, based on a previous characterization that showed significantly higher expression levels and inducibility for this region compared to truncated versions^35^. A TAA stop codon was appended to terminate the N-terminal region of the OsmY protein contained in this region and a strong RBS^22^ was inserted immediately after the stop codon to form the stress sensor promoter which we call pOsmY (Supp. Fig. 17). The actuator protein GRP was discovered when the N-terminal region of the OsmY protein was instead fused directly to the N-terminus of GFP using a Ser-Ala-Gly linker. Although the mechanism of action is not essential for our findings, the growth-modulating effect of GRP is most likely due to the signaling peptide (residues 1 to 28) that usually targets OsmY to the periplasm^36^.

A complete list of plasmids and strains can be found in Supp. Fig. 17 and Supplementary Information 1, respectively.

### Culture Media and Inducers

All plate reader and microscopy experiments were performed in EZ rich media (Teknova) prepared according to the manufacturer’s instructions with 0.2% glucose as the carbon source, except for the gating fidelity tests shown in Supp. Fig. 13d and e which were conducted in lysogeny broth (Difco). To prevent cell adhesion to microfluidic channels, all media were supplemented with 0.075% (wt/vol) Tween 20 (for consistency also in plate reader experiments). Selection antibiotics (50*μ*g/ml Ampicillin, 50*μ*g/ml Kanamycin and/or 34*μ*g/ml Chloramphenicol) and basal inducers were added as required. A stock suspension of 1 *μ*m-diameter fluorescent beads (Polysciences, Fluoresbrite Multifluorescent, Cat no. 24062, visible in RFP channel) was created at a ratio of one drop per 250*μ*l of MilliQ water. The resulting suspension was added to all microfluidic media at a dilution of 1:1000 to be able to measure flow velocity in the media channels (see Microfluidics and Microscopy section). The specific AHL used in this study is 3-oxo-C6-homoserine lactone, also known as Auto Inducer 1 for *Vibrio fischeri.* Large volumes of chemicals/inducers (for example, 5M NaCl in MilliQ water used to reach a target concentration of 500mM) were added by replacing the corresponding amount of water in the EZ rich recipe.

### Microfluidics and Microscopy

Microfluidic experiments were conducted using the techniques established in previous studies from our lab^18,37^. Briefly, PDMS was poured on a silicon wafer mold (created using photolithography) and baked at 80°C for one hour before cutting individual chips, punching fluidic access ports, and cleaning bonding surfaces. PDMS pieces containing single chips were then bonded to standard glass cover slips after 3min exposure in a Jelight UV-Ozone cleaner and kept overnight at 80 °C for strong bonding. A schematic of the specific microfluidic device used in this study can be found in Supp. Fig. 1.

The night before the experiment’ a culture of the corresponding strain was inoculated from a −80 °C glycerol stock into lysogeny broth (Difco) supplemented with the appropriate selection antibiotics (see Supplementary Information 1 for a list of strains) and grown overnight in a shaking incubator at 37 °C. On the day of the experiment, the saturated overnight culture was diluted 1:1000 into 5ml of the same media supplemented with 0.075% (wt/vol) Tween 20 and incubated until an OD600 of 0.1 was reached. 1ml of this culture was spun down at 3000rcf in a standard tabletop microcentrifuge for 3min and the cells were resuspended in 5*μ*l of the base media used for the microfluidic experiment (see Culture Media section). The cell suspension was then immediately loaded into the waste port of the microfluidic device, which had been degassed in a vacuum chamber for at least 30min and base media was loaded into all other ports. After traps were seeded’ media and waste ports were connected to the corresponding reservoirs and media channels were flushed with base media at high flow rate (>2000*μ*m/s) before calibrating the flow rate to 100*μ*m/s in the media channels by measuring streak length of fluorescent beads at 1s exposure in the RFP imaging channel. A Nikon Eclipse Ti2 inverted microscope was used with a Photometrics CoolSNAP HQ2 CCD camera to capture images and a Lumencor SOLA SE LED light engine for fluorescent excitation. The stage holding the microfluidic device was enclosed in a plexiglass incubation chamber maintaining a constant 37°C environment. The acquisition software was Nikon Elements®.

We performed time-lapse imaging with two settings: At a magnification of 10x, we used exposure times of 500 *μ*s for phase contrast, 500 ms for GFP (25% LED intensity) and 500 ms for RFP (20% LED intensity). All 14 traps for a single media composition (7 for each of the two sizes) could be captured in a single image, requiring 8 images for the whole chip, taken every 4min. High-resolution data (e.g., for analysis of growth patterns as in Figs. 1b, 1c) was acquired at 30x, with exposure times of 20 ms for phase contrast, 100 ms for GFP (25% LED intensity) and 500 ms for RFP (20% LED intensity). With one image for each trap size and media composition (16 images), each containing two traps, an imaging interval of 2 min was possible.

### Microscopy Data Analysis

Image registration using custom MATLAB™ scripts was performed on phase-contrast images to correct for stage jitter. Masks for individual traps and media channels were automatically generated for data extraction. Dye intensity measured in media channels was used to calculate media compositions generated by the microfluidic mixer (Supp. Figs. 1, 2). We analyzed the growth pattern in the traps by calculating inter-image differences (IIDs) between consecutive phase-contrast images in order to detect cell movement. To account for subpixel residual jitter after registration, IIDs between individual pixel intensities were averaged over a 5×5 pixel area before taking the absolute value. We created kymographs by averaging intensities from the appropriate imaging channels (or IIDs for growth) along the horizontal direction of the trap for each time point and concatenating the resulting columns chronologically.

### Plate Reader Experiments

Microplate experiments were conducted in standard flat-bottom 96-well plates (Falcon) in a Tecan Infinite M200 Pro plate reader at 37 °C. GFP fluorescence was recorded with an excitation wavelength of 485 nm and an emission wavelength of 520 nm, RFP fluorescence was recorded at 580 nm / 620 nm. Both fluorescence and 600 nm-absorbance (OD600) were recorded every 3.5 min. Experiments were started from a 1:1000 dilution of saturated cultures. Lysis strains were preincubated for 1 h before adding inducer to prevent growth delays due to priming with LuxR at the previous growth arrest. A background signal obtained from the first 10 measurements in each well was subtracted from all OD and fluorescence traces. Fluorescence was then divided by OD600 to yield a normalized fluorescence signal.

The gating fidelity for the stress-gated RFP circuit (Fig. 3c, Supp. Figs. 13d, e) was calculated as the ratio between the peak normalized RFP fluorescence signal for the well and the normalized fluorescence at OD 0.7. (Supp. Fig. 13a). Lysis efficiency was defined as the percentage reduction in OD 1 h after reaching the peak OD. Exponential cell doubling rates (Fig. 3f, Supp. Fig. 14) were calculated as the slope of the least-squares linear fit to log_2_[OD(t)] between OD600 of 0.08 and 0.2.

### Numerical Simulations

Chemical reaction kinetics of the sensor and actuator components and population dynamics in batch culture as well as dynamics of these components and nutrients in spatially extended colonies were simulated using custom scripts written in MATLAB™. For the spatiotemporal model, we constructed a one-dimensional reaction-diffusion-advection model using a split-step finite-difference implicit-diffusion numerical scheme. A detailed description of the model equations, assumptions and parameters can be found in the Supplementary Information 3.

### Data availability

The data that support the findings of this study are available from the corresponding authors upon request. Addgene catalog numbers for plasmids will be provided upon publication.

### Code availability

The modeling code for the numerical simulations is available from the corresponding authors upon request.

