## Supplementary Figures and Movie Captions for "Engineered phenotype patterns in microbial populations"

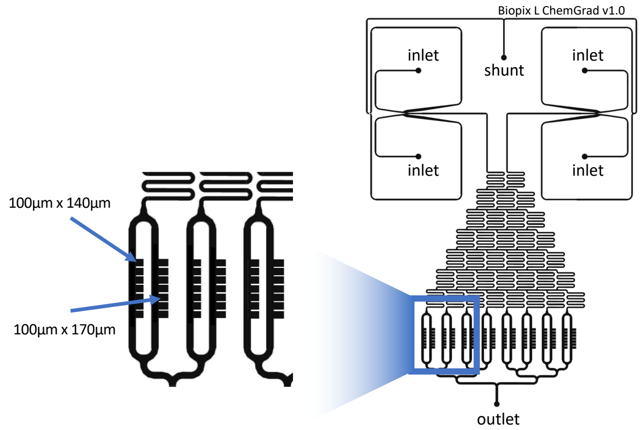


**Supplementary Figure 1 Microfluidic device used in this study.** The device features 4 inlets, a shunt preventing backflow into inlet ports by receiving currently unused media, and an outlet. Inlet hydrostatic pressure can be varied by changing media reservoir heights to select the top or bottom inlet on either side of the chip. For time-dependent induction experiments, typically, only the shunt, one port on the left side, two ports on the right side and the outlet were punched to create the sockets for plugging in reservoir connections. The three inlet ports are fed from reservoirs containing the growth media, where one reservoir on the right side additionally contains the inducer. A linear mixer creates 8 different mixtures – from 100% left growth medium to 100% right growth medium – which are each fed into two channels, to which the cell traps are attached (two different depths, 7 traps each: $100\mu m \times 140\mu m$ and $100\mu m \times 170\mu m$). The height (distance from glass cover slip to PDMS ceiling) is 1.6 to $1.8\mu m$ within the cell traps and $40\mu m$ for the channels, similar to the "biopixels" design used in previous studies from our group[18]. Media at different inlets contain different amounts of fluorescent dye to retrospectively determine the inducer concentration from microscopy images (see Supp. Fig. 2).

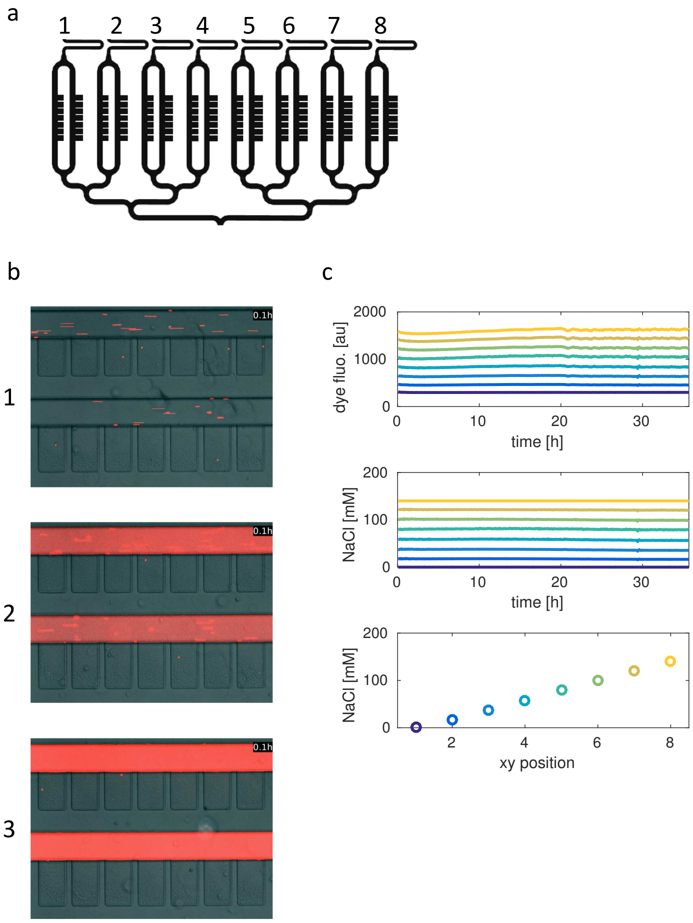


**Supplementary Figure 2 On-chip mixing of two different growth media. a**, Trapping region of the microfluidic chips with numbered media supply channels containing different media mixtures. **b**, Example microscopy images of trapping regions connected to media supply channels 1, 2 and 3. One of the two media fed into the mixer contains a red fluorescent dye, leading to distinct fluorescence intensities in the different channels. The small streaks also visible in the images are fluorescent beads used to determine flow rate. **c**, (*Top*) Averaged raw fluorescence signal measured over time in each of the 8 imaging positions (i.e. channel regions with different media). Colors indicate channel number from blue (position 1) to yellow (position 8). (*Middle*) Raw fluorescence values are then normalized by minimum and maximum fluorescence at each time point. Assuming that the highest fluorescence corresponds to 100% of media *with* dye and the lowest fluorescence corresponds to 100% media *without* dye, differential media composition is calculated for each position by linear scaling between these min and max values at each time point (in this case, the media with dye contained 140mM NaCl). The purity of media in the two channels yielding min and max fluorescence is verified throughout each experiment by imaging additional positions upstream of the trapping region. (*Bottom*) Average calculated chemical composition at each position over the entire time of the experiment.

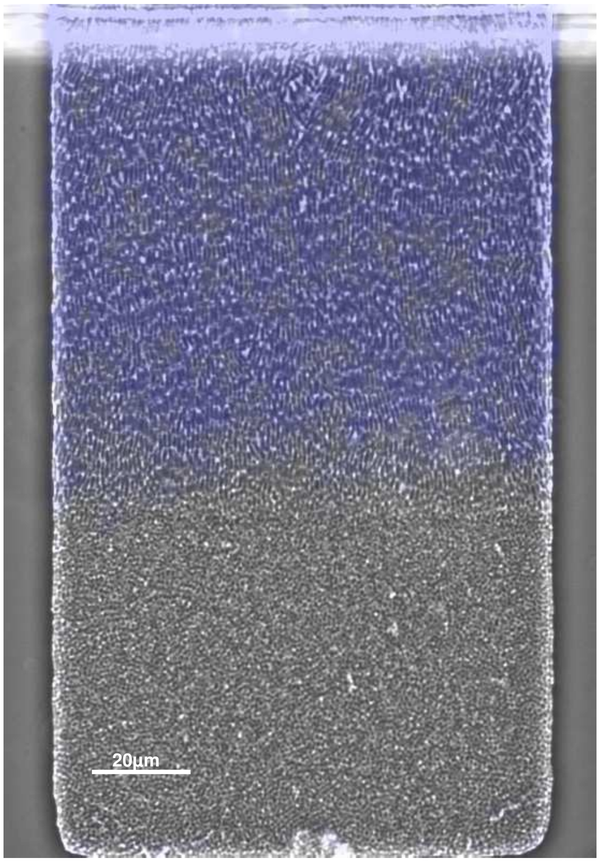


**Supplementary Figure 3  High-resolution image of full microfluidic trap after establishment of phenotypic heterogeneity.**

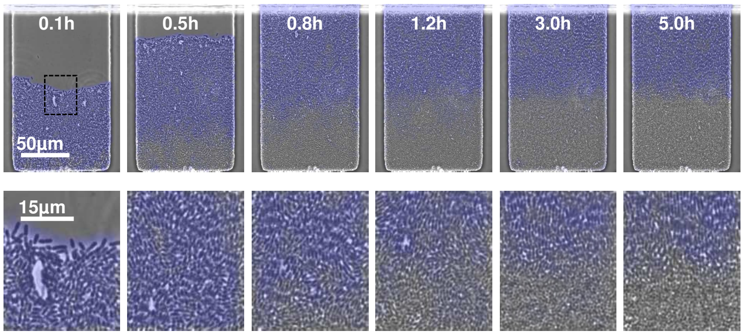


**Supplementary Figure 4  Initial growth in the microfluidic trap.** Time-lapse images as in Fig. 1b with zoomed-in region around the eventual growth boundary (see also Supplementary Movie 1).

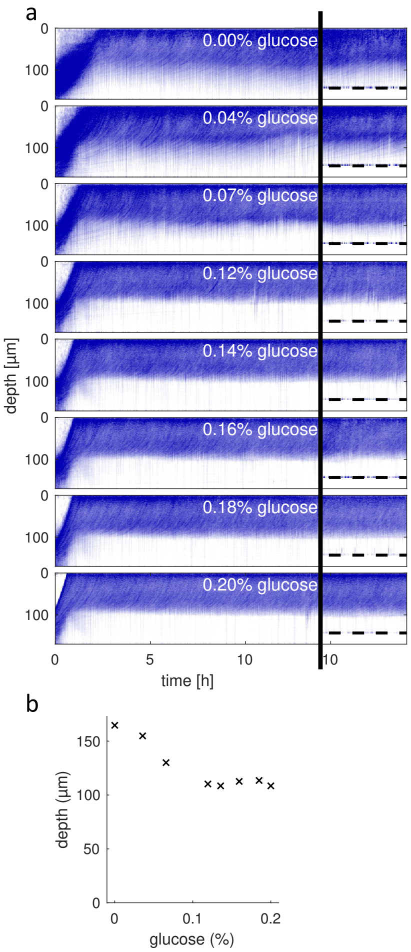


**Supplementary Figure 5 Growth patterns in different media compositions.** **a**, Kymographs as in Fig. 1c during the establishment of a steady state growth pattern in 170$\mu$m deep traps for growth media containing different amounts of glucose (the standard concentration is 0.2% w/v). For lower concentrations, growth extends further into the back of the trap with a smoother transition between regions of growth and no growth. Kymographs on the right of the black line show consistent behavior in the smaller (140$\mu$m deep) traps (the dashed line indicates the end of the trap). **b**, Distance from the mouth of the microfluidic trap at which the inter-image difference (IID) shown in panel a drops below detectable levels. For this purpose, the IID profile was averaged over 4h of steady-state growth. The threshold for growth detection was chosen as the lowest possible level that safely detected the no-growth region in standard growth medium.

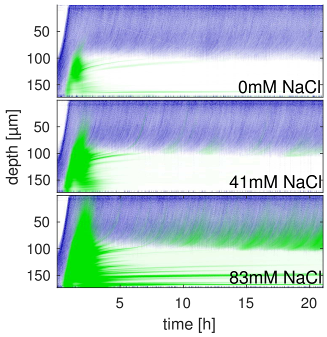


**Supplementary Figure 6 Stress sensor activation at different basal stress levels.** Kymographs showing growth (blue) and GFP fluorescence (green) for lower NaCl concentrations. Compared to Fig. 1f, the fluorescence signal has been rescaled to visualize even weak activation of the pOsmY stress sensor. While the sensor is transiently activated upon growth arrest even in the absence of basal osmotic stress (0mM NaCl), higher NaCl also elicits a measurable response in dividing cells close to the growth boundary.

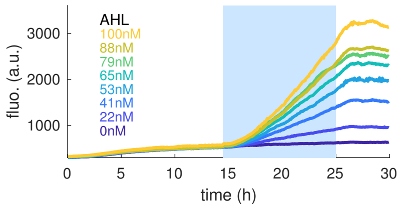


**Supplementary Figure 7 Expression in growth-arrested cells.** Expression of untagged RFP from the pLuxI promoter. As in Figs. 1g and h, the plot shows average fluorescence measured in the growth-arrested back of the microfluidic trap. The blue shaded area marks the window of induction with AHL.

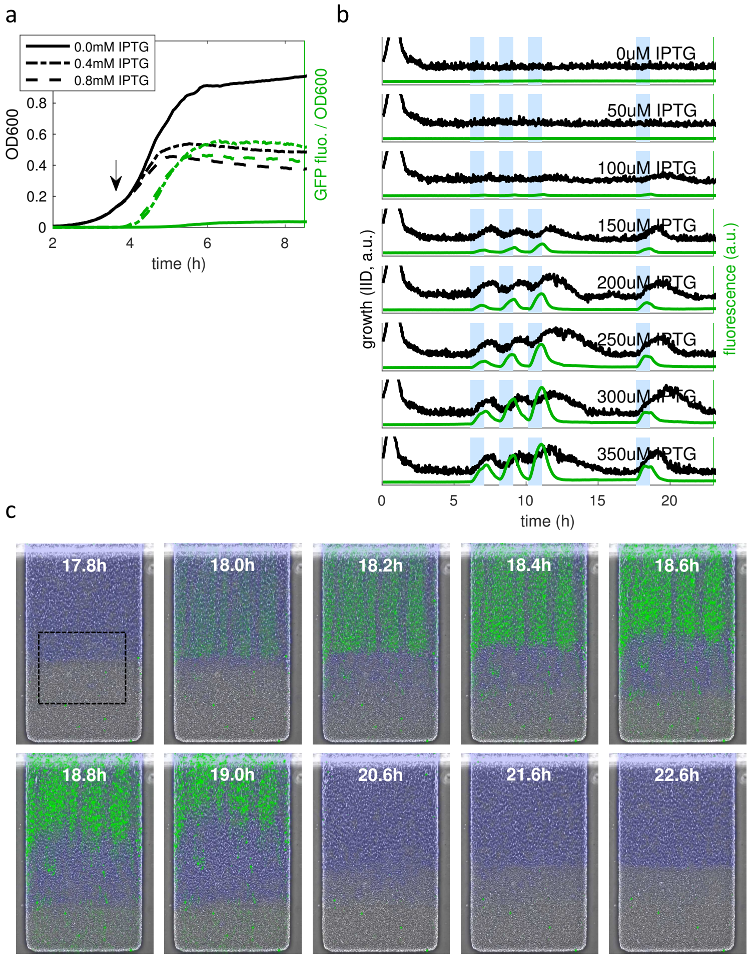


**Supplementary Figure 8 Actuator circuit in batch culture and microfluidics.** **a**, IPTG was added when the culture of *E. coli* cells equipped with the actuator circuit reached an OD600 of 0.13. Growth is halted through expression of GRP-ssrA from pLlacO-1 at a higher OD compared to Fig. 2a, consistent with the later addition of IPTG. **b**, Time series of growth (calculated as inter-image difference, IID, see Methods) and fluorescence in our microfluidic environment, averaged over the area in the trap indicated in panel c. Blue shaded regions mark the induction windows for all 8 IPTG concentrations tested. Cumulative effects can be observed when pulses of IPTG are given in short succession. **c**, Time-lapse images of the experiment shown in panel b during the last pulse of 300$\mu$M IPTG (see Supplementary Movie 3). The pulse begins at t = 17.6h.

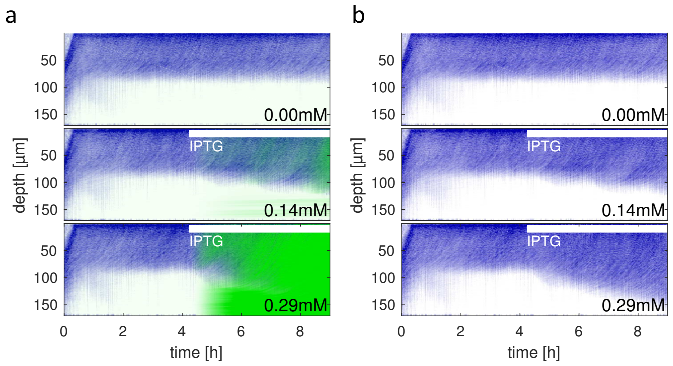


**Supplementary Figure 9 Continuous induction of actuator circuit.** **a**, Kymographs of cell populations carrying the actuator circuit, which is induced continuously starting around t = 4h. With increasing GRP-ssrA expression, the area of active growth extends further and further into the trap. **b**, Same data as in panel a, omitting fluorescence for better visualization of the growth pattern.

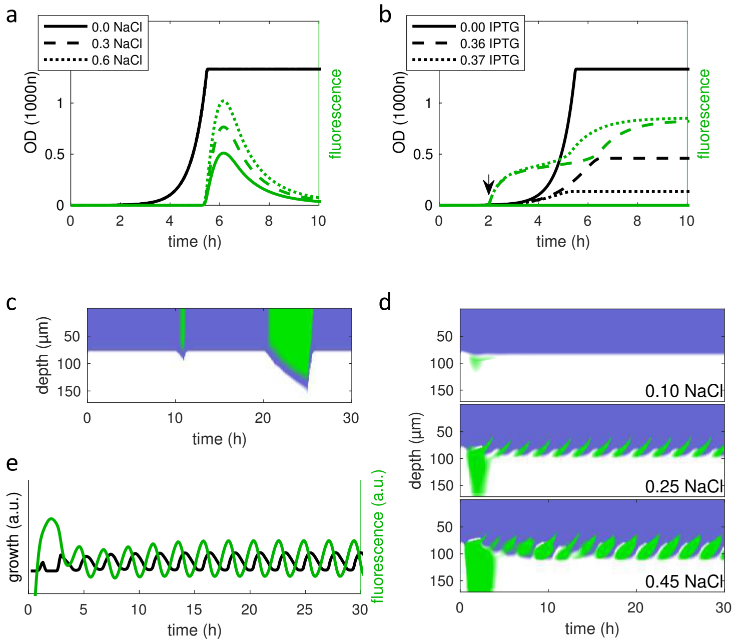


**Supplementary Figure 10 Numerical simulations of the sensor-actuator circuit.** **a**, Activation of the stress sensor upon growth arrest in numerical simulations of a well-mixed batch culture of cells. The promoter strength of the sensor is modeled to increase with the NaCl concentration in the media (cf. Fig. 1e). **b**, Actuator and population dynamics in batch culture upon IPTG induction. The arrow marks the start of induction. The production rate of the actuator protein is proportional to the IPTG induction level, halting growth earlier for higher IPTG (cf. Fig. 2a). **c**, Spatiotemporal simulations of actuator protein induction and population dynamics in a microfluidic trap. 1h and 5h pulses of IPTG (amplitudes 0.5) lead to reversible growth resumption in the growth arrested region of the simulated microcolony (cf. Fig. 2b and Supp. Fig. 9). **d**, Numerical simulations of the full spatiotemporal model of the sensor-actuator circuit coupled with population and nutrient dynamics. Oscillations near the growth interface are observed for sufficiently high induction (NaCl) levels of the sensor promoter. **e**, In the oscillatory layer of the microcolony (panel d), induction of the actuator protein is followed by growth resumption, which is followed by growth arrest and induction of the actuator protein from the stress sensor, restarting the cycle (cf. Fig. 2g).

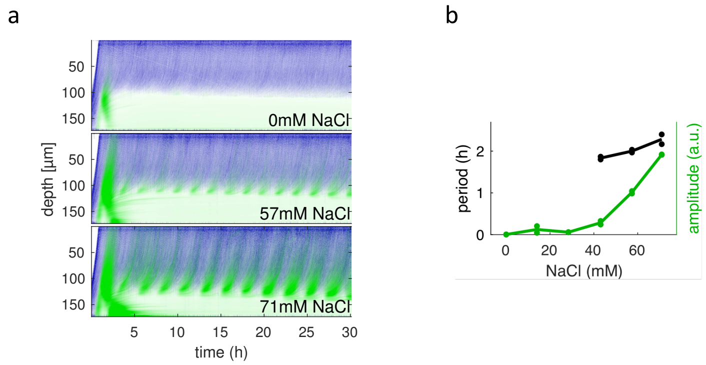


**Supplementary Figure 11 Induction of oscillations in the sensor-actuator circuit.** **a**, Kymographs of populations carrying the sensor-actuator circuit (Fig. 2c). Basal osmotic stress (NaCl) activates the sensor close to the interface (Fig. 1f) and initiates negative feedback leading to oscillations. **b**, Period and amplitude of fluorescence oscillations. Period is only shown for NaCl concentrations yielding finite oscillation amplitudes. Data points for each concentration correspond to two traps imaged at high resolution (see Methods).

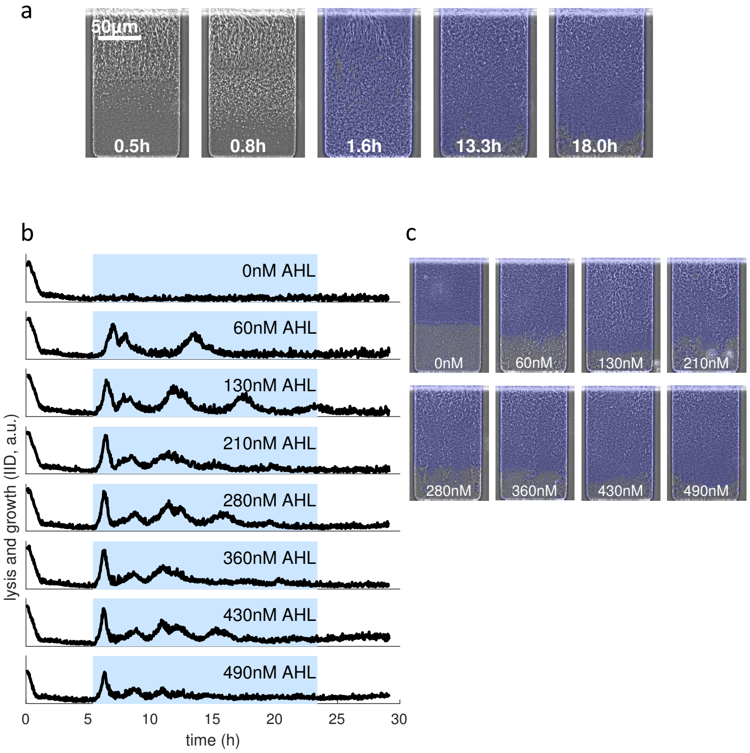


**Supplementary Figure 12 Unsuccessful elimination of growth-arrested cells with the ungated lysis circuit. a**, Time-lapse images of unsuccessful phenotype elimination with the ungated lysis circuit (LuxR-pLuxR-pLuxI-E-ssrA-AAV) for induction with 490nM AHL (cf. panels b and c). Lysis starts in the growing part of the population, causing growth-arrested cells to receive fresh nutrients and resume growth, while simultaneously also lysing. Cells regrowing despite circuit activation form disorganized colonies and display no visible phenotype pruning. Time stamps denote time after beginning of induction. See also Supplementary Movie 5. **b**, Growth and lysis dynamics (calculated via Inter-image differences, see Methods) in the region of the trap usually occupied by growth-arrested cells for different AHL induction levels of the ungated lysis circuit. After the initial lysis, no sharp lysis events are observed. Instead, uncoordinated phases of spatially inhomogeneous lysis, regrowth and growth arrest without lysis lead to fluctuating IID that settles at a low level (cf. end states in panel c). **c**, Images taken after 18h of AHL induction with the indicated concentration, when traps have settled into a steady state. While cells in all traps initially lyse (cf. panel b), no distinct phenotype elimination is observed and colony structure is disorganized (see Supplementary Information 2).

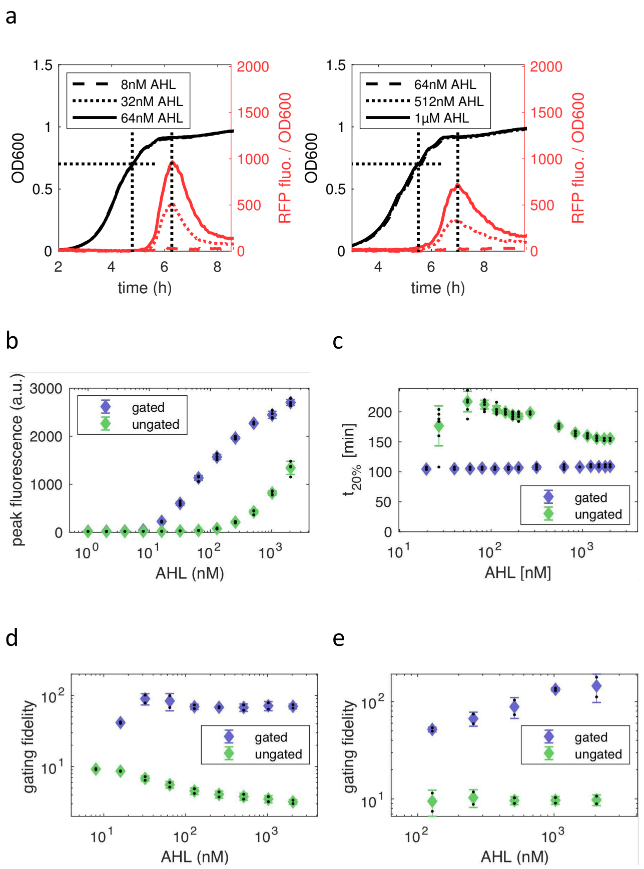


**Supplementary Figure 13 Stress-gated expression of RFP. a**, Example traces of OD and RFP fluorescence in plate reader experiments with the gating circuit (Fig. 3a) with RFP-ssrA-AAV as the target gene (*left*), compared to ungated expression from a standard bidirectional LuxR-pLuxR-pLuxI cassette with constitutive LuxR (*right*). Plotted induction levels were chosen to show similar levels of expression. **b**, Peak fluorescence in plate reader experiments of the same RFP-ssrA-AAV gating circuit (Fig. 3a). The stress-gated circuit exhibits increased sensitivity at low concentrations, which enabled us to determine gating fidelity (Fig. 3c) for much lower concentrations for this circuit compared to constitutive LuxR. Error bars represent SD across 4 technical replicates. **c**, We tested the same circuits in microfluidics and measured the fluorescence in the growth-arrested region in the back of the trap in response to a 10h induction with different AHL concentrations. $t_{20\%}$ is the time until 20% of peak fluorescence (as shown in Fig. 3d) are reached. Error bars represent SD across 7 traps. **d**, Gating fidelity for stress-gated and ungated RFP-ssrA-AAV from plate reader experiments as in Fig. 3c, but measured in LB. Error bars represent SD across 2 technical replicates. **e**, Gating fidelity for stress-gated and ungated RFP-ssrA-AAV from plate reader experiments as in Fig. 3c, but measured in LB supplemented with 0.2% glucose. Error bars represent SD across 2 technical replicates.

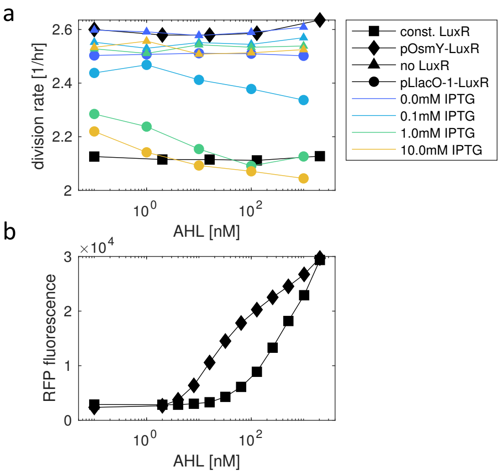


**Supplementary Figure 14 Expression of LuxR impacts growth rate during exponential growth.** **a**, We compared growth rates in the plate reader between OD 0.08 and 0.2 for cells expressing untagged RFP from the AHL-inducible pLuxI promoter. 4 different strains were tested that differed in their expression of the LuxR regulator protein: (1) constitutive LuxR from the traditional bidirectional LuxR-pLuxR-pLuxI cassette; (2) pOsmY-driven LuxR as in our gating circuit (Fig. 3a, where target gene = RFP-ssrA); (3) no LuxR at all; (4) LuxR expressed from the IPTG-inducible pLlacO-1 promoter. The constitutive and the gating circuit show consistently low and high growth rates, respectively, across all AHL concentrations. The pLlacO-1-LuxR construct approaches the same low growth rates upon induction with IPTG, whereas the no-LuxR control shows no such effect, suggesting that it is indeed LuxR that causes slowed growth. **b**, Peak RFP fluorescence of the constitutive-LuxR circuit and and the gating circuit from panel a. Expression from the gating circuit is significantly increased for lower concentrations of AHL, indicating sufficient expression of LuxR at later growth phases.

a

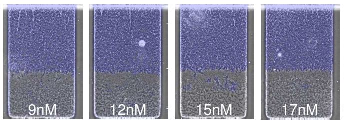


b

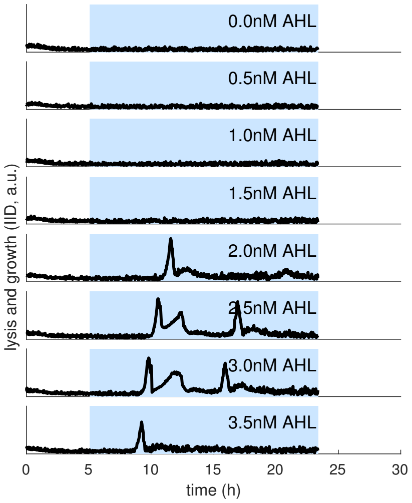


**Supplementary Figure 15 Behavior of the SGLC at low AHL concentrations near onset. a**, Images taken after 24h of AHL induction with the indicated concentration, when traps have settled into a steady state. Lower concentrations do not settle into steady states (see Supp. Fig. 16) or show imperfect pruning. **b**, Growth and lysis dynamics (calculated via Inter-image differences, see Methods) in the region of the trap usually occupied by growth-arrested cells. No lysis occurred below AHL concentration of 2nM. For higher concentrations and lysis onset times across parameters, see Fig. 4d and e, respectively.

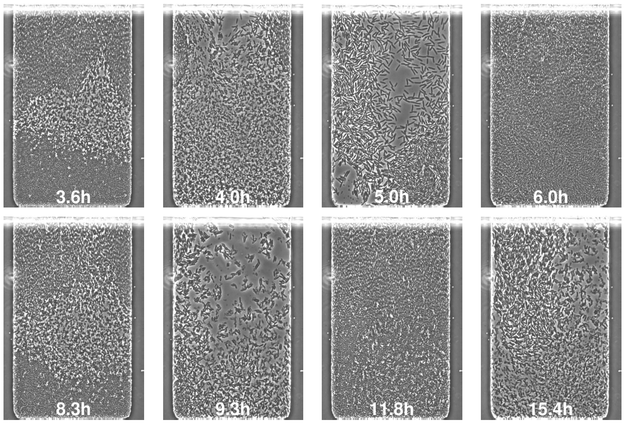


**Supplementary Figure 16 Oscillatory lysis.** Time-lapse images showing oscillatory behavior of the SGLC close to onset of lysis (3nM AHL, cf. Fig. 4d and Supp. Fig. 15b). Long delay between LuxR priming by the gating circuit and accumulation of sufficient lysis protein (Fig. 4e) causes sequential fill-up and lysis (see Supplementary Movie 7). Time stamps denote time after beginning of induction.

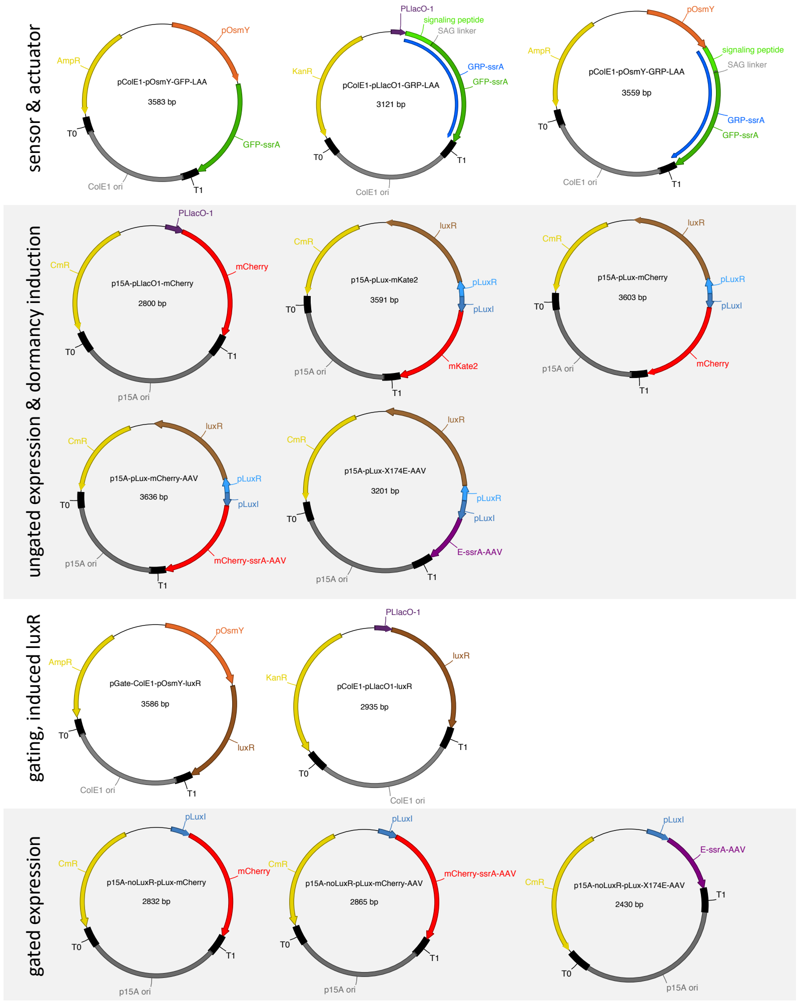


**Supplementary Figure 17 Maps of all plasmids used in this study.** "ssrA" in annotations and "LAA" in plasmid names both refer to the strong ClpXP-specific ssrA degradation tag with amino-acid sequence "AANDENYALAA". "ssrA-AAV" and "AAV" refer to the weaker degradation tag "AANDENYAAAV"[34]. For details regarding assembly and strains, see Methods and Supplementary Information 1.

**Supplementary Movie 1 Establishment of phenotypic heterogeneity in 170$\mu$m deep traps.** The data in this movie corresponds to Fig. 1b and Supp. Fig. 4.

**Supplementary Movie 2 pOsmY sensor activation during pattern formation and close to the no-growth boundary.** The data in this movie corresponds to the time-lapse images in Fig. 1f.

**Supplementary Movie 3 Actuator induction modulates the growth pattern.** The data in this movie corresponds to the time-lapse images in Supp. Fig. 8c.

**Supplementary Movie 4 A diffusion-mediated spatiotemporal feedback loop leads to oscillations in growth and fluorescence.** The data in this movie corresponds to the time-lapse images in Fig. 2f.

**Supplementary Movie 5 Unsuccessful elimination of dormant cells by the ungated lysis circuit.** The data in this movie corresponds to the time-lapse images in Supp. Fig. 12a.

**Supplementary Movie 6 The stress-gated lysis circuit (SGLC) as a phenotype filter.** The data in this movie corresponds to the time-lapse images in Fig. 4a.

**Supplementary Movie 7 Oscillatory lysis with the SGLC at low AHL concentrations.** The data in this movie corresponds to the time-lapse images in Supp. Fig. 16.
