## Supplementary Information for "Engineered phenotype patterns in microbial populations"

### Supplementary Information
### Engineered phenotype patterns in microbial populations

Philip Bittihn,[1,2,3] Andriy Didovyk,[1,4] Lev S. Tsimring,[1,2,†] and Jeff Hasty[1,2,5,6,†]

*[1]BioCircuits Institute, University of California, San Diego, La Jolla, California, USA*
*[2]The San Diego Center for Systems Biology, La Jolla, California, USA*
*[3]Current address: Max Planck Institute for Dynamics and Self-Organization, Göttingen, Germany*
*[4]Current address: Vertex Pharmaceuticals, San Diego, California, USA*
*[5]Department of Bioengineering, University of California, San Diego, La Jolla, California, USA*
*[6]Molecular Biology Section, Division of Biological Sciences, University of California, San Diego, La Jolla, California, USA*

†

#### 1 Plasmids and strains

All strains in this study were created by transforming the host strain MG1655Z1 with plasmids of the ColE1 and p15A family. MG1655Z1 expresses high levels of the repressors TetR and LacI, the latter of which was required for induction experiments with the pLlac-O1 promoter in non-dividing cells (Fig. 1g) as well as the controlled LuxR expression experiments (Supp. Fig. 14a). Therefore, it was used for all experiments for consistency. The following table lists the strains, ordered by their first appearance, along with the contained plasmids, selection antibiotics (ampicillin: Amp, chloramphenicol: Cm, kanamycin: Kan) and references to figures containing experimental data obtained from them. Maps for all constructs can be found in Supp. Fig. 17. Plasmid names use the following conventions: "LAA" and "AAV" refer to the strong and weak ssrA degradation tags with amino acid sequences "AANDENYALAA" and "AANDENYAAAV", respectively.[1] "pLux" refers to the standard AHL-inducible expression cassette with divergent expression of LuxR and the protein of interest (cf. Supp. Fig. 17). Constructs missing the LuxR portion of the cassette are labeled "noLuxR" explicitly. The gating plasmid (bold) enables gated expression from such noLuxR constructs (cf. Fig. 3a).

| Strain | Plasmids | Figs. | Supp. Figs. |
|---|---|---|---|
| PB01 | p15A-pLux-mKate2 (Cm) | 1b, 1c | 3, 4, 5 |
| PB02 | pColE1-pOsmY-GFP-LAA (Amp)<br>p15A-pLux-mKate2 (Cm) | 1f, 1e | 6 |
| PB03 | p15A-pLlacO1-mCherry (Cm) | 1g | |
| PB04 | p15A-pLux-mCherry-AAV (Cm) | 1h, 3c, 3d, 3f | 13 |
| PB05 | pColE1-pLlacO1-GRP-LAA (Kan) | 2a, 2b | 8, 9 |
| PB06 | pColE1-pOsmY-GRP-LAA (Amp)<br>p15A-pLux-mKate2 (Cm) | 2f, 2g, 2h | 7, 11 |
| PB07 | p15A-pLux-X174E-AAV (Cm) | 3e-h | 12 |
| PB08 | p**Gate**-ColE1-pOsmY-luxR (Amp)<br>p15A-noLuxR-pLux-mCherry-AAV (Cm) | 3c, 3d, 3f | 13 |
| PB09 | p15A-noLuxR-pLux-mCherry (Cm) | | 14 |
| PB10 | pColE1-pLlacO1-luxR (Kan)<br>p15A-noLuxR-pLux-mCherry (Cm) | | 14 |
| PB11 | p15A-pLux-mCherry (Cm) | | 14 |
| PB12 | p**Gate**-ColE1-pOsmY-luxR (Amp)<br>p15A-noLuxR-pLux-mCherry (Cm) | | 14 |
| PB13 | p**Gate**-ColE1-pOsmY-luxR (Amp)<br>p15A-noLuxR-pLux-X174E-AAV (Cm) | 3e-h, 4 | 15, 16 |

#### 2 Unsuccessful phenotype elimination using standard pLuxI promoter

Our first attempt to eliminate the growth-arrested phenotype from the population, before the construction of the gating module, was to take advantage of the non-dividing cells' capacity for protein expression and degradation (Figs. 1g, h) using our standard, bidirectional, AHL-inducible pLuxI expression cassette (identical to that used for RFP-ssrA; Fig. 1h), as it appeared to support strong expression and had a natural preference for the dormant cells. In this cassette, LuxR is constitutively expressed from its native pLuxR promoter (Supp. Fig. 17). However, when we placed ssrA-AAV-tagged lysis protein E under the control of this standard pLuxI expression cassette and the construct was activated in the microcolonies with sufficient AHL, dividing cells began lysing first, causing dormant cells to start regrowing before also lysing (Supp. Fig. 12a; Supplementary Movie 5). After regrowth, the colonies eventually settled into a disorganized state of constant partial lysis for all effective AHL concentrations (Supp. Fig. 12b, c), without any visible phenotype elimination. The fact that growth in these final steady-states could be observed *deeper* in the trap than without lysis (compare different end states to 0 nM in Supp. Fig. 12c), indicated that the expression of the lysis protein continuously affects the growth pattern of *dividing* cells, either creating "holes" or slowing down growth, and thus allowing nutrients to diffuse deeper into the trap. In addition, we still observed dormant cells in the back of the trap, seemingly unaffected by lysis. These could be descendants of previously dividing cells in which mutations deactivated the lysis module due to the immense selective pressure. Alternatively, they indicate insufficient expression of lysis protein. All of these problems seemed to point to one common root cause: the expression level of of lysis protein E from the standard pLuxI promoter was too high in dividing cells compared to growth-arrested cells. This motivated the construction of the gating module that would shift expression towards the non-dividing phenotype.

#### 3 Numerical Modeling

Our model of the full sensor-actuator circuit describes the coupled dynamics of the cellular concentration of the growth repression *protein* p expressed from the sensor promoter, and the *capacity* c of the cell to activate the stress-responsive sensor promoter. In a single cell, the model equations are

$$\dot{p} = P(f,p,c) - g(f,p) \cdot p \tag{1a}$$

$$\dot{c} = C(f,p,c) - g(f,p) \cdot c \tag{1b}$$

with

$$P(f,p,c) = \beta \cdot c \cdot \exp\Big(-F_1 \cdot (f - f_{crit})^2\Big) - \gamma_p \cdot p \tag{2}$$

$$C(f,p,c) = g(f,p) - \gamma_c \cdot c. \tag{3}$$

denoting synthesis and degradation rates for the corresponding components that depend on their own concentrations and the concentration of the critical nutrient concentration (“food") f. By choosing sufficiently large $F_1 \gg 1$ we enforce that the growth repression protein (GRP) is expressed from the stress-activated sensor promoter only when f is near a specific (low) value of $f = f_{crit}$ (“metabolic stress”) proportional to the cell’s stress response capacity c. The synthesis rate is also modulated by the effective promoter strength of the sensor $\beta$, which can be increased by additional NaCl in the medium: $\beta = 0.6 + \mathrm{NaCl}$ (Fig. 1e). As the proteins that we express from the sensor promoter are ssrA-tagged and ClpXP is active in stationary phase in batch as well as non-growing cells in microfluidics (cf. Figs. 1e and 1h), p has an intrinsic degradation rate $\gamma_p$.

In addition to synthesis and degradation, both p and c are also diluted with a rate that is equal to the cellular growth rate $g(f,p)$, reflected by the second terms in the r.h.s. of both Eqs. (1). The stress response capacity c represents the ability of the cell to elicit a stress response due to food scarcity. At any given instant, it tends towards a value of $g(f,p)/[\gamma_c + g(f,p)]$). Its nominal (maximum) value in growing cells is therefore $g_0/(\gamma_c + g_0)$. When a cell stops growing ($g(f,p) = 0$), its level of c decays exponentially with rate $\gamma_c$ towards zero. After growth resumption, the capacity c first has to recover sufficiently for cells to activate the sensor promoter in response to nutrient stress, as suggested by the lack of an immediate stress response in non-dividing cells close to the no-growth boundary (Supp. Fig. 6) and cells that have just resumed growth (Fig. 2f).

The growth rate itself depends on the nutrient concentration f and the amount of GRP p,

$$g(f,p) = g_0 \cdot \max\{0, \tanh[F_2 \cdot (f - f_{crit})]\} \cdot \max\Big\{0, 1 - (p/p_{crit})^3\Big\}, \tag{4}$$

where $g_0$ is the maximum growth rate. The first max{...} term on the right hand side models the sharp (we choose $F_2 \gg 1$) growth rate dependence on the food concentration near the critical food concentration $f_{crit}$ where it drops to zero. The second max{...} term models the reduction in growth rate by the growth repression protein, reaching 0 for a critical value $p_{crit}$.

The dynamics of the nutrient concentration f depends on the specific experimental setup: For example, in batch culture, it is depleted from initial concentration $f_0$ with the rate proportional to the cell density n and the per-cell nutrient consumption rate $F(f,p)$

$$\dot{f} = n \cdot F(f,p) = -n \cdot \max\{\alpha \cdot g(f,p), \alpha_0\} \tag{5}$$

This expression indicates that f is consumed with a rate that is proportional to the growth rate g, but never falls below a basal metabolic turnover rate $\alpha_0$ even if cells do not grow.

Equations (1) – (5) represent the basal model of the sensor-actuator circuit.

##### Decoupling sensor and actuator components

We decouple the sensor (promoter) and actuator (growth repression protein, GRP) components in the above model in order to compare them separately to the experiments of the isolated sensor and actuator modules. Modeling of the sensor module (Fig. 1d) is straightforward, as all the induction characteristics of p are preserved. However, since p in this case is simply GFP-ssrA (without the growth-modulating effect of GRP), we exclude its effect on the growth rate and replace Eq. (4) by

$$g^*(f) = g_0 \cdot \max\{0, \tanh(F_2 \cdot (f - f_{crit}))\}. \tag{6}$$

We then interpret p simply as the fluorescence without any feedback on cell growth.

To model the isolated actuator module (with IPTG-controlled induction of GRP, Fig. 2a), we replace Eq. (2) in Eq. (1a) for the growth repression protein p with

$$P^*(f,p,c) = P^*(f,p) = IPTG \cdot \max\{\delta, \tanh(F_2 \cdot (f - f_{crit}))\} - \gamma_p \cdot p, \tag{7}$$

which decouples it from the cell's stress response (c is irrelevant in this context). In this case, we assume that the synthesis of GRP is proportional to the concentration of IPTG and the instantaneous growth rate $g^*(f)$, however below $f_{crit}$ it remains small but finite ($\delta \ll 1$), in accordance with microfluidic experiments that show a much-weakened response in non-growing cells, even though dilution is completely absent (Fig. 2b). The residual production rate is required to account for the fact that GRP levels remain high even in stationary phase despite being ssrA-tagged (Fig. 2a) and that for high enough induction, expression can eventually be detected in non-growing cells (cf. Fig. 2b and Supp. Fig. 9).

##### Batch culture model

In batch culture, the cell density n grows with the instantaneous growth rate $g(f,p)$:

$$\dot{n} = g(f,p) \cdot n \tag{8}$$

To model the dynamics of the sensor module in the batch culture, we integrate the basal model, Eqs. (1), (5), together with this equation for the cell density, where we replace g with $g^*$ from Eq. (6). Numerical simulations lead to the temporal dynamics shown in Supp. Fig. 10a. Similarly, to model the batch culture experiment with the isolated actuator module induced by IPTG, we integrate the basal model and the cell density equation with P replaced by $P^*$, Eq. (7). Supplementary Figure 10b shows the results of this simulation for three values of the IPTG concentration. Both of these numerical simulations closely match experimental observations shown in Figs. 1e and Fig. 2a.

#### Spatiotemporal model

To model the populations of cells harboring these modules in the spatial context of a microfluidic trap, we construct a one-dimensional reaction-diffusion-advection model that generalizes the intracellular basal model (1) and the food equation (5), where p, c and f become fields over a spatial coordinate x. $x = 0$ corresponds to the mouth of the trap and $x = L$ corresponds to the back wall. For simplicity, we assume a constant cell density throughout the trap, eliminating the need for an explicit dynamic variable n (we set $n = n_{trap}$ everywhere, see "Parameters" section for details). We impose Dirichlet boundary condition $f(0) = f_0$ at $x = 0$ to model fresh media supply at the mouth of the trap, and no-flux boundary conditions for all variables at $x = L$. Extracellular $f(x,t)$ is only subject to diffusion with diffusion constant $D_f$, whereas $p(x,t)$ and $c(x,t)$ are intracellular quantities which are not diffused, but advected through the trap due to cell motion caused by growth. Assuming that cells in the microfluidic trap are densely packed and incompressible, the velocity $v(x,t)$ is determined by the instantaneous, cumulative cell growth rate in the trap

$$v(x,t) = \int_L^x g(f(x,t), p(x,t))\,dx, \tag{9}$$

(the velocity is always zero at the back wall $x = L$). Note that this velocity field is *not* divergence-free due to local growth. This also means that dilution of p and c is already implicitly modeled through advection (which is consistent since the divergence of the velocity field $\frac{\partial v}{\partial x}$ is exactly $g(f,p)$). Hence, the full system of partial differential equations is

$$\frac{\partial p}{\partial t} = P(f,p,c) - \frac{\partial (pv)}{\partial x} \tag{10a}$$

$$\frac{\partial c}{\partial t} = C(f,p,c) - \frac{\partial (cv)}{\partial x} \tag{10b}$$

$$\frac{\partial f}{\partial t} = n_{trap} \cdot F(f,p) + D_f \frac{\partial^2 f}{\partial x^2} \tag{10c}$$

We simulated this system of advection-reaction-diffusion equations using implicit finite difference first-order in time and second-order in space integrator in MATLAB™ using the same parameter values as for batch culture simulations. Numerical simulations of this system with the actuator modification, Eq. (7), lead to Supp. Fig. 10c, reproducing the experimental observation of growth resumption in Fig. 2b and Supp. Fig. 9. Simulations of the full system of equations without modifications produced Figs. 2d, e and Supp. Figs. 10d, e.

#### Parameters and initial conditions

To make sure that our batch culture and spatiotemporal simulations are consistent with each other, and that the physical scales of growth and nutrient consumption/transport are realistic, we chose them accroding to the following strategy: The growth rate was taken directly from plate reader experiments (cf. Fig. 14). For the cell density n, we assumed natural, dimensionless units of the fractional volume occupied by cells. In these units, an OD600 of 1 corresponds to approximately $n = 0.1\%$ or $10^{-3}$ according to the literature[2]. We arbitrarily chose $f_0 = 1$ as the initial nutrient concentration in fresh media and assumed that the culture starts from $n(t = 0) = 10^{-7}$. The metabolite consumption rate $\alpha$ was then adjusted such that the culture reached saturation (i.e. f reaches $f_{crit}$)

when the OD is of order 1 (because of exponential growth, the value of $f_{crit}$ itself only has a weak influence as long as $f_{crit}$ is significantly smaller than $f_0$).

In the context of the microfluidic trap, cells are densely packed, such that, in the same dimensionless units, $n_{trap}$ (the constant density assumed throughout the trap) should be of order 1. To account for the rod shape of the cells and non-optimal packing, we chose a value of $n_{trap} = 1/3$. Diffusion coefficients of metabolites in aqueous solutions range between 500 and 2000 $\mu m^2 s^{-1}$, [3, pp. 32, Table 2.1], including potentially relevant aminoacids[4]. In light of the crowded environment in the trap, we chose a value at the lower end of this range, $D_f = 500\,\mu m^2 s^{-1}$, consistent with a recent study on biofilm front propagation[5]. Without further adjustments, the combination of diffusion, cell density and metabolite consumption rates (from the batch culture setting) indeed caused the metabolite concentration to reach $f_{crit}$ within the boundaries of a $L = 170\,\mu m$-deep traps (Supp. Fig. 10d) leading to a growth pattern that closely resembles the experimental observation. We therefore concluded that the physical scales of our two modeling settings were consistent.

It is interesting to note that the parameters $\alpha$, $\alpha_0$ and $D_f$ are in a regime where the time scale of Eq. (10c) is sufficiently short for the diffusion process to essentially become a quasi steady state: At any given time point, the profile of f is very close to the steady state of Eq. (10c), and, therefore, increasing all three parameters further by a common factor does not alter model dynamics. Effects only start being noticed when these parameters are reduced by about two orders of magnitude. Thus, while diffusion provides the feedback for the oscillator circuit (Fig. 2c), it does not cause any significant delay by itself, as communication through diffusion is virtually instantaneous.

Other rates were chosen to closely match experimental observations for the sensor and actuator components. Note that $\beta$ simply scales p, and p's effect on growth rate, Eq. (4), is unaltered if $p_{crit}$ is adjusted accordingly, so only the ratio $\beta/p_{crit}$ (or IPTG/$p_{crit}$ for externally induced GRP) is a real model parameter.

The following table lists the parameters used for all numerical simulations in this study. Concentrations are dimensionless.

| parameter | value |
|---|---|
| $L$ | $170\,\mu m$ |
| $D_f$ | $18 \times 10^5\,\mu m^2 h^{-1} = 500\,\mu m^2 s^{-1}$ |
| $n_{trap}$ | 1/3 |
| $f_0$ | 1 |
| $g_0$ | $2.5 \cdot \log(2)\,h^{-1}$ |
| $\alpha$ | $600\,h^{-1}$ |
| $\alpha_0$ | $120\,h^{-1}$ |
| $f_{crit}$ | 0.2 |
| $F_1$ | 75 |
| $F_2$ | 50 |
| $\delta$ | 0.01 |
| $\beta$ | $(0.6 + \text{NaCl})\,h^{-1}$ |
| $p_{crit}$ | 0.4 |
| $\gamma_p$ | $1\,h^{-1}$ |
| $\gamma_c$ | $0.3\,h^{-1}$ |

Initial conditions at $t = 0$ were $f = f_0$ (also used as the Dirichlet boundary condition at $x = 0$ in the spatial case), $p = 0$ and $c = g_0/(\gamma_c + g_0)$. In batch culture, we assumed an initial cell density (vol/vol) of $n = 10^{-7}$ (i.e. roughly OD 0.0001). As described in the previous section, in the spatiotemporal model, we assume a constant cell density throughout the trap of $n = n_{trap} = 1/3$.

1. Andersen, J. B. *et al.* New Unstable Variants of Green Fluorescent Protein for Studies of Transient Gene Expression in Bacteria. *Applied and Environmental Microbiology* **64**, 2240–2246 (1998).

2. Sezonov, G., Joseleau-Petit, D. & D'Ari, R. Escherichia coli physiology in Luria-Bertani broth. *Journal of Bacteriology* **189**, 8746–8749 (2007).

3. Stein, W. D. *Channels, Carriers, and Pumps*. An Introduction to Membrane Transport (Academic Press, 1990).

4. Ma, Y., Zhu, C., Ma, P. & Yu, K. T. Studies on the Diffusion Coefficients of Amino Acids in Aqueous Solutions. *Journal of Chemical & Engineering Data* **50**, 1192–1196 (2005).

5. Wang, X., Stone, H. A. & Golestanian, R. Shape of the growing front of biofilms. *New Journal of Physics* **19**, 125007 (2017).
